# Parallel systems for sound processing and functional connectivity among layer 5 and 6 auditory corticothalamic neurons

**DOI:** 10.1101/447276

**Authors:** Ross S. Williamson, Daniel B. Polley

## Abstract

Cortical layers (L) 5 and 6 are populated by a spatially intermingled menagerie of neurons with distinct inputs and downstream targets. Here, we made optogenetically guided recordings from L5 and L6 corticothalamic (CT) neurons in the mouse auditory cortex to discern underlying patterns of functional connectivity and sensory processing in the largest sub-cerebral projection system. Whereas L5 CT neurons showed broad stimulus selectivity with sluggish response latencies and extended temporal non-linearities, L6 CTs exhibited sparse sound feature selectivity and rapid temporal processing. L5 CT spikes lagged behind neighboring units and imposed weak feedforward excitation within the local column. By contrast, L6 CT spikes drove robust and sustained activity in neighboring units. Our findings underscore a duality among CT projection neurons, where L5 CT units are canonical broadcast neurons that integrate sensory inputs for transmission to distributed downstream targets, while L6 CT neurons are positioned to regulate thalamocortical response gain and selectivity.

## Introduction

Corticofugal neurons fall into two distinct classes: intratelencephalic and sub-cerebral (Harris & Shepherd 2015; Harris & Mrsic-Flogel 2013). In the auditory cortex (ACtx), intratelencephalic projection neurons are often found in layer (L) 5a, project locally to L2/3 and distally to ipsi-and contralateral cortex and striatum (Games & Winer 1988; Winer & Prieto 2001; Yuan et al. 2011). By contrast, L5b and L6 ACtx sub-cerebral projection neurons target auditory processing centers in the brainstem, midbrain and thalamus, as well as numerous targets outside of the central auditory pathway, including the striatum and lateral amygdala (Winer 2006; Bajo & King 2013; Asokan et al. 2018).

The largest compartment of the corticofugal projection system comes from neurons in layer 5 and 6 of the cortex that innervate the thalamus (Ojima 1994; Prieto & Winer 1999; Winer et al. 2001). L5 corticothalamic (CT) axons deposit giant terminal collaterals in the dorsal division of the medial geniculate body (MGBd, (Bajo et al. 1995; Rouiller & Welker 2000;Ojima 1994)), en route to additional downstream targets in the tectum, striatum, and amygdala (Veinante et al. 2000; Bourassa et al. 1995; Bourassa & Deschênes 1995; Deschênes et al. 1986;Rockland 1998; Kita & Kita 2012; Asokan et al. 2018). By contrast, the sub-cerebral projections of L6 CT neurons are restricted to the thalamus, dropping dense deposits of axon collaterals among GABAergic neurons in the thalamic reticular nucleus and terminating most extensively in the ventral subdivision of the MGB (MGBv, (Lund et al. 1979; Staiger et al. 1996; Chevée et al. 2018; Llano & Sherman 2008; Cai et al. 2018; Guo et al. 2017; Zhang & Jones 2004)). Unlike L5 CT neurons, L6 CT axons also collateralize vertically within the local cortical column, predominantly synapsing onto neurons in layers 5a and 6 (Guo et al. 2017; Olsen et al. 2012;Kim et al. 2014; Prieto & Winer 1999).

The distinct patterns of connectivity among L5 and L6 CT neurons hint at their potential parallel roles in sensory processing and cortical feedback. L5 CT neurons, with their far-ranging axons and elaborate dendritic processes are regarded as the canonical “broadcast” neurons of the cortex, pooling inputs from the upper layers and transmitting signals to widespread downstream targets. L6 CT neurons, by contrast, appear specialized for thalamocortical gain control given that their axons terminate in all three nodes of the thalamo-cortico-thalamic loop: the MGB, GABAergic cells in the thalamic reticular nucleus and intrinsically, within the local column. Differences in anatomical connectivity of L5 and L6 CT neurons are paralleled by additional dichotomies in cell morphology, intrinsic membrane properties, and synaptic properties in these cell types (Ojima 1994; Diamond et al. 1969; Bajo et al. 1995; Rouiller & Welker 2000; Sherman & Guillery 2011; Llano & Sherman 2009; Bartlett et al. 2000; Andersen et al. 1980; Sherman 2016; Joshi et al. 2016; Dembrow et al. 2010).

Until recently, technical limitations have made it difficult to extend observations made in fixed tissue or acute brain slice to the realm of sensory processing in awake animals. Here, we leveraged advances in multi-channel electrophysiology and optogenetics to make targeted recordings of ACtx CT neurons in awake mice. We show that L5 CT neurons utilize dense, nonlinear coding of sound features and have little influence on local cortical processing whereas L6 CT neurons have sparse selectivity for sound features, and strongly modulate local processing within ACtx. These findings indicate that each class of CT projection neuron performs distinct operations on incoming sensory signals, which likely impart distinct effects on their downstream targets.

## Results

### Two types of corticothalamic projection neurons

As a first step towards highlighting the differences in L5 and L6 CT neurons, we first wanted to confirm that the well-established patterns of sub-cerebral connectivity in other brain areas and species could be reprised in the mouse auditory system. To visualize L5 CT neurons, we used an intersectional virus strategy that exploited their divergent axons fields that also innervate the tectum (Asokan et al. 2018). We first injected canine adenovirus 2 (CAV2-Cre) into the inferior colliculus, which selectively infects local axon terminals to express retrogradely in upstream projection neurons (N=6 mice, **Figure 1A**: *left*) (Soudais et al. 2001; Schneider et al. 2014; Liu et al. 2016). We then made a second injection of a Cre-dependent virus in the ipsilateral ACtx to label the corticotectal axon fields as well as other potential downstream targets (Figure 1A: *left).* As expected, we observed a smattering of smaller neurons that project to the midbrain in L6b, but a majority cells in L5b with prominent apical dendrites (Figure 1A: *left*) (Schofield 2009). In addition to dense terminal labeling in the external and dorsal cortex of the IC (**Figure 1C**: *left*), we also observed terminal labelling in the MGBd (**Figure 1B**: *left*). These findings confirmed previous reports that corticotectal projection neurons could also be described as CT projection neurons, with the caveat that we do not know what fraction of L5 corticotectal cells are also corticothalamic (Bajo et al. 1995; Rouiller & Welker 2000).

We used the Ntsr1-Cre transgenic mouse to label L6 CT neurons based on our prior report that 97% of Ntsr1-positive neurons in the ACtx are L6 CT, and 90% of all L6 CT neurons are Ntsr1-positive (Guo et al. 2017). We injected a Cre-dependent virus into the right ACtx, allowing a fluorescent reporter to be expressed throughout L6 CT axon fields (Figure 1A: *right*). As expected, we observed labelled cell bodies in layer 6a with a band of neuropil labeling in L5a (Figure 1A: *right*) and a dense plexus of axon terminal labelling in the MGB, predominantly in the ventral subdivision (Figure 1D: *right*). Labelling was not observed in the ipsilateral IC (Figure 1C: *right*), indicating that the Ntsr1-Cre expressing cells are separate from other classes of L6 projection neurons that have more distributed local and subcerebral targets (Briggs 2010; Sundberg & Granseth 2018).

**Figure 1.**
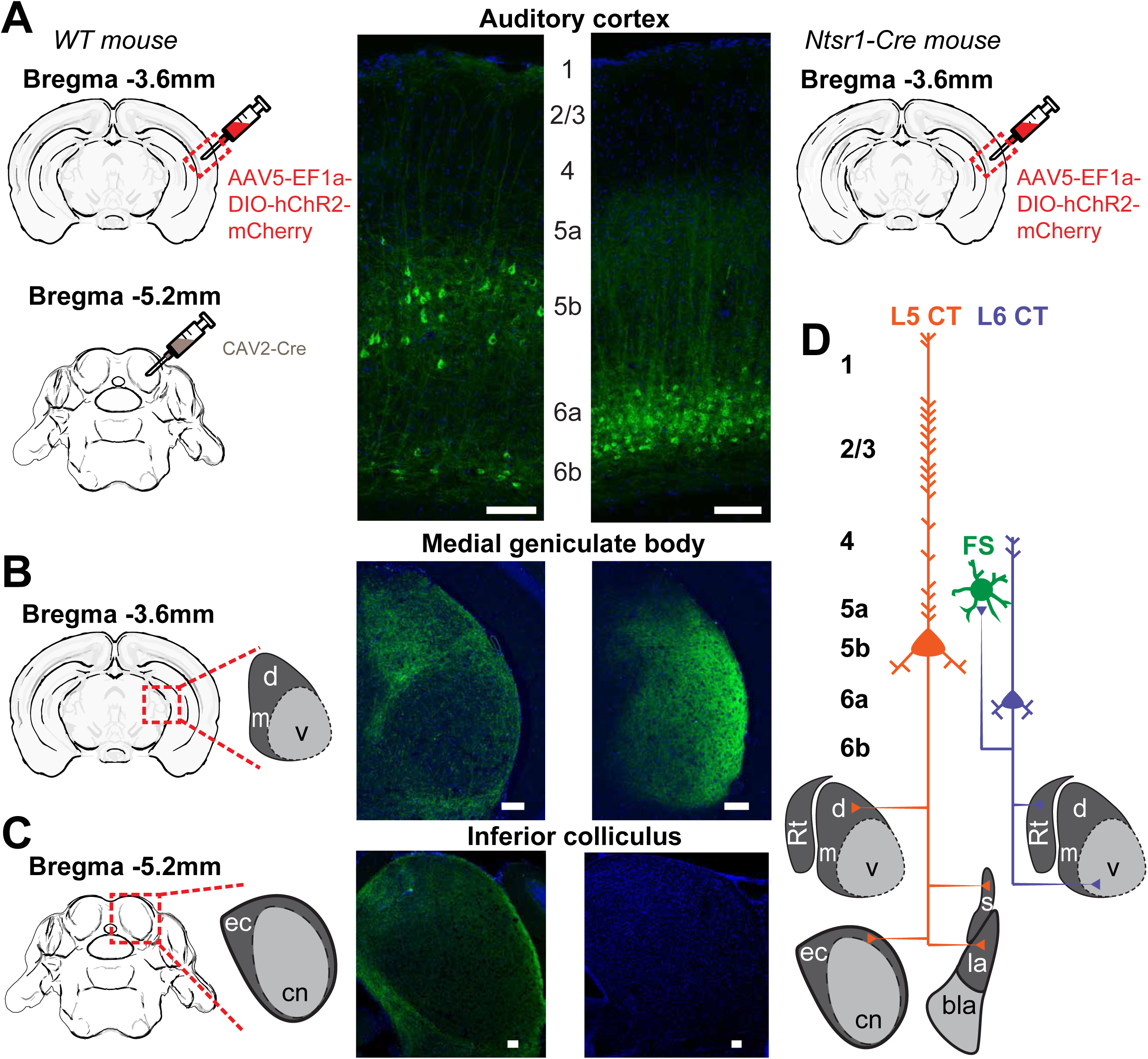
Dual corticothalamic pathways from L5 and L6. **A**: Illustration of transgenic and viral strategy used to selectively label L5 neurons that project to the MGB and inferior colliculus (left) or L6 neurons that project to the MGB (right). Cell body locations and intracortical processes for CAV2-Cre- and Ntsr1-Cre-expressing ACtx neurons. **B**: 20x confocal sections of the ipsilateral MGB in both L5 CT (*left*) and L6 CT (*right*) mice. The L5 projection shows strong axon labelling in the MGBd, while the L6 projection shows strong axon labelling in the MGBv. **C**: 20x confocal sections of the ipsilateral IC in both L5 CT (*left*) and L6 CT (*right*). The L5 projection shows strong axon labelling in the external cortex of IC, while the L6 projection shows none. **D**: Schematic of dual corticothalamic projection pathways from the ACtx. Abbreviations: d = MGBd, m = MGBm, v = MGBv, Rt = Thalamic Reticular Nucleus, ec = External Cortex, cn = Central Nucleus, s = Striatum, la = Lateral Amygdala, bla = Basolateral Amygdala. Scale bars = 100 μm.

### Antidromic phototagging of ACtx projection neurons

To make targeted recordings from L6 CT neurons, we expressed ChR2 in the ACtx of Ntsr1-Cre mice and implanted an optic fiber such that the tip was positioned lateral to the MGB (**Figure2A**). Because the intersectional labeling experiment demonstrated that L5 CT neurons projecting to the MGBd also innervate the inferior colliculus, we targeted L5 CT neurons by implanting an optic fiber atop the dorsal cap of the inferior colliculus of WT mice (**Figure 2B**). We made extracellular recordings from awake, head-fixed mice (N=12) using high-density 32- channel silicon “edge” probes that primarily spanned L5 and 6 (**Figure 2C**). Spikes were sorted into single-unit clusters (n=1,246) using Kilosort (Pachitariu et al. 2016). We used the characteristic pattern of laminar current sinks and sources to assign each recorded neuron to L5 (n=509) or L6 (n=625, **Extended Figure 2A-B**) (Kaur et al. 2005; Guo et al. 2017). The peak-to-trough delay of each unit’s waveform was used to classify cells as either regular-spiking (RS) or fast-spiking (FS) (RS: n=1,028, FS: n=184, **Extended Figure 2C**).

**Figure 2.**
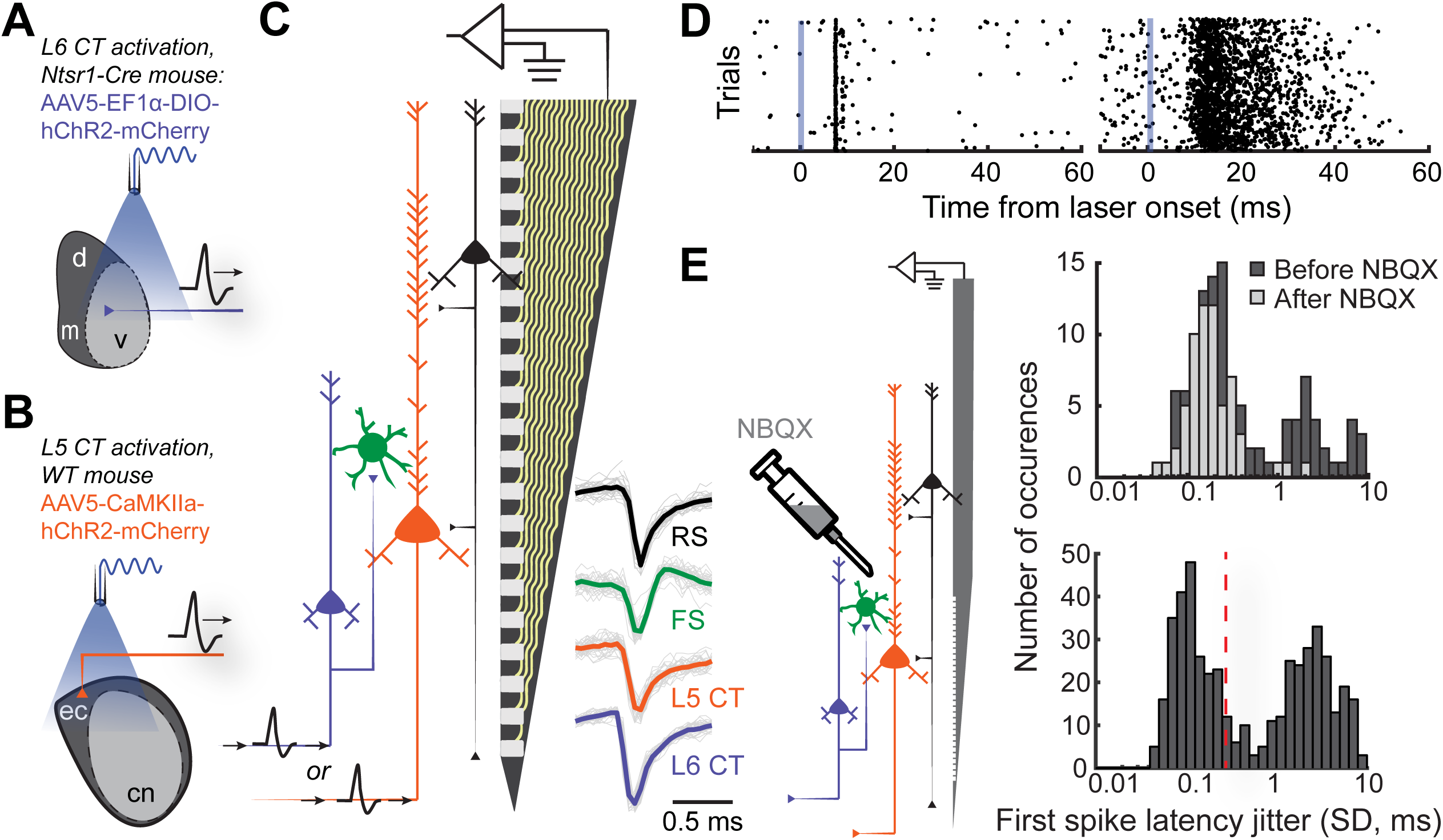
Antidromic phototagging of auditory cortical projection neurons. **A**: Illustration of the strategy to activate neurons in ACtx L6 that project to the MGB (L6 CT). **B**: Illustration of optogenetic approach to isolate L5 ACtx neurons with axons that innervate the inferior colliculus and MGB (L5 CT). **C**: Schematic of the 32-channel silicon edge probe with 20 μm inter-contact spacing used for extracellular recording, alongside four major neuron classes. Example mean spike waveforms (thick lines) are shown with individual waveforms (thin lines). **D**: Example unit spike rasters of low- and high-jitter antidromic activity (*left* and *right,* respectively). **E**: Illustration of the strategy used to block local synaptic transmission with NBQX (*left*). First spike latency jitter histograms from anesthetized pharmacology experiments (top), and the awake dataset showing the 0.3 SD cutoff for identifying directly activated cells (*bottom*).

To identify L5 and L6 CT projection neurons, we optogenetically activated their axon terminals and documented the temporal patterning of antidromic action potentials (**Figure 2D**). Consistent with previous observations, we hypothesized that antidromically activated spikes with very low trial-to-trial jitter in arrival time were non-synaptic events reflecting direct photoactivation, whereas high-jitter spikes likely arose from intracortical synaptic transmission within the ACtx (Li et al. 2015; Jennings et al. 2013; Lima et al. 2009). To verify this, we blocked local glutamatergic transmission within ACtx with local application of NBQX, an AMPA receptor antagonist (**Figure 2E**) in a subset of anesthetized control mice (N=4, n=188). NBQX did not affect the low-jitter mode of the distribution but eliminated spikes with variable first spike latencies, confirming that the low-jitter mode of the distribution reflected direct activation of the recorded projection neuron, whereas higher jitter spikes arose through local polysynaptic activation (Figure 2E: *top*). Based on our pharmacology control experiments, we operationally identified recordings of antidromically phototagged L5 and L6 CT neurons in our awake recordings as spikes with laser-evoked first spike latency jitter of 0.3 ms SD or less (Figure 2E: *bottom;* L5 CT: n=132, L6 CT: n=83). As further evidence of this, we estimated the laminar locations of the phototagged populations and confirmed that they matched the expected anatomy (**Extended Figure 2D**). Few of our phototagged CT neurons had spike shapes in the FS range (L5 CT: n=6, L6 CT: n=8), suggesting that these projection neurons are largely separate from the recently described population of parvalbumin-expressing GABA projection neurons (Zurita et al. 2018; Rock et al. 2017).

### Sensory characterization of CT projections

To characterize sensory processing differences in optogenetically identified L5 and L6 CT units, we first quantified pure tone frequency response areas (FRAs). Only neurons that had well-defined FRAs were included for analysis (L5: n=174, L6: n=172, L5 CT: n=44, L6 CT: n=27). From each FRA, we computed the tuning bandwidth (20 dB above threshold, **Figure 3A**) and the tone-evoked onset latency (**Figure 3B**). While there were no overall differences in tuning bandwidth between neurons in L5 and 6, L5 CT neurons were more broadly tuned than L6 CT neurons (Wilcoxon Rank Sum test, p=0.03, **Figure 3C**). Neurons in L5, including the L5 CTs, exhibited tone-evoked first spike latencies that were approximately twice as long as L6 (Wilcoxon Rank Sum test; layer: p<2×10^-9^, cell-type: p<4×10^-4^, **Figure 3D**).

**Figure 3.**
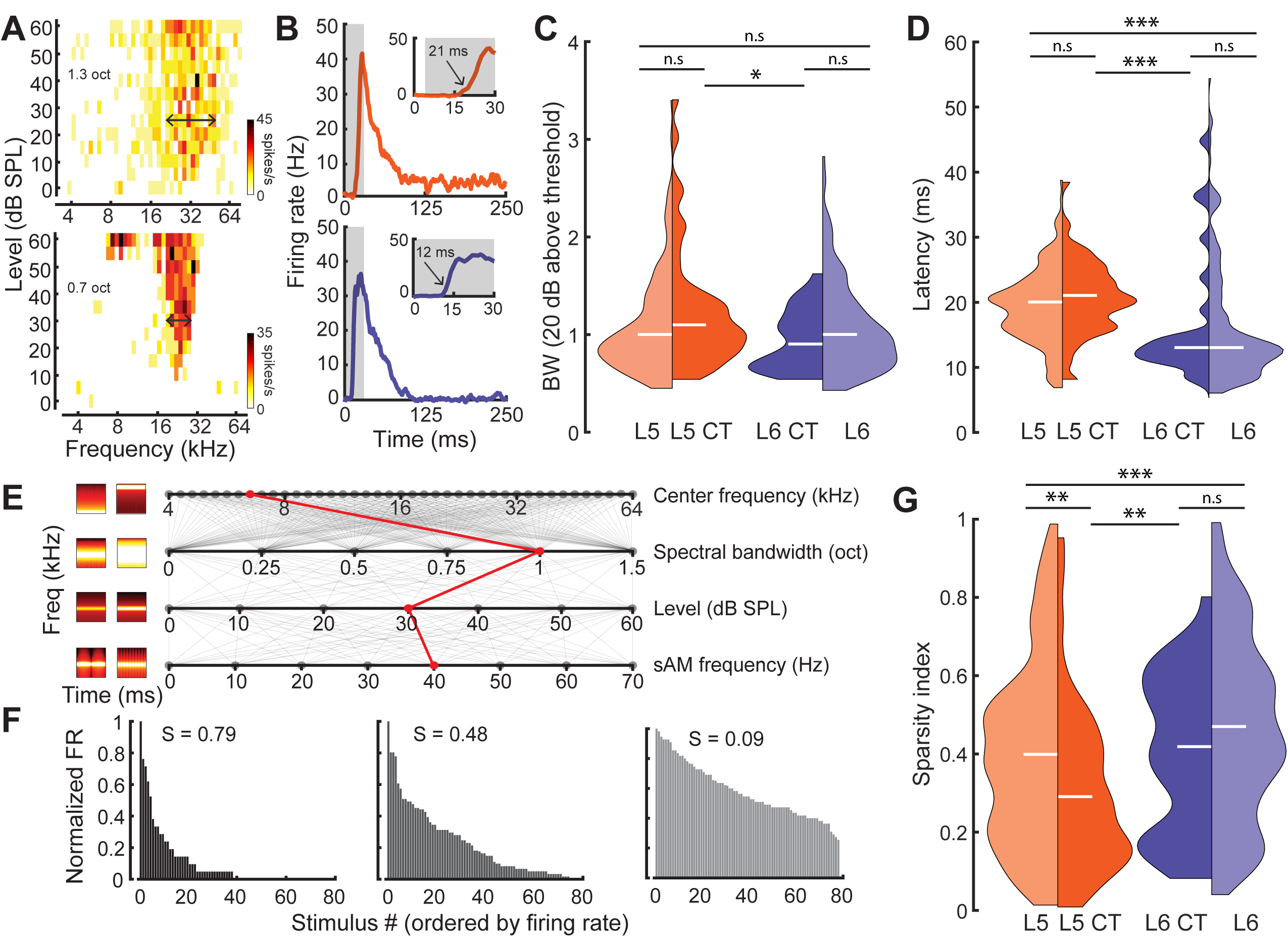
Sensory characterization of CT projections. **A**: Example FRAs from a broadly-tuned L5 CT neuron(*top*) and a narrowly tuned L6 CT neuron (*bottom*). **B**: Example PSTHs from a long-latency L5 CT neuron (*top*) and a short-latency L6 CT neuron (*bottom*). **C**: Split-violin plots showing the FRA bandwidth distributions. White horizontal line represents the median. **D**: Tone-evoked latency distributions for the 4 different groups. **E**: Spectrotemporally modulated noise burst tokens varied across four acoustic dimensions: Center frequency, spectral bandwidth, level, and sinusoidal amplitude modulation (sAM) frequency, represented here from top to bottom. Frequency x time stimulus spectrograms depict a lower (*left*) and higher (*right*) value for each corresponding acoustic parameter. Red line indicates an example of stimulus values used for a single trial. **F**: Example histograms for firing rate across the same 80 randomly selected spectrotemporal noise burst tokens. High sparsity values (S) indicate a narrow distribution with enhanced stimulus selectivity (*left*) while low sparsity values indicate a broad distribution with reduced stimulus selectivity (*right*). **G**: Split-violin plots showing the sparsity distributions for the 4 different groups. Asterisks denote statistically significant differences at the following levels: *p<0.05, **p<0.01, ***p<0.005, as determined by the Wilcoxon Rank Sum test.

As a next step, we generated a set of spectrotemporally modulated noise bursts, that varied in center frequency (4-64 kHz, 0.1 octave steps), spectral bandwidth (0-1.5 octaves, 0.1 octave steps), level (0-60 dB SPL, 10 dB SPL steps), and sinusoidal amplitude modulation rate (0-70 Hz, 10 Hz steps) (**Figure 3E**). We then used a standard measure of sparsity to quantify the shape of each cell’s response distribution (**Figure 3F**) (Rolls & Tovee 1995; Vinje & Gallant 2000; Chambers et al. 2014). This lifetime sparseness index is bounded between 0 and 1, with values close to 1 reflecting selectivity for a sparse set of stimuli (Figure 3F: *left*) and values close to 0 reflecting a broad response distribution (Figure 3F: *right*). Responses in L6 were sparser than L5 (Wilcoxon Rank Sum test, p<3×10^-3^, **Figure 3G**) and complex sound selectivity was sparser in L6 CTs than L5 CTs, in agreement with the differences in pure tone tuning bandwidth (Wilcoxon Rank Sum test, p=0.02, Figure 3G). Tuning bandwidth and sparsity were inversely correlated (Pearson’s r=-0.3, p<2×10^-8^, **Extended Figure 3A**), but we found no other significant correlations between other tuning parameters (sparsity vs latency: Pearson’s r=0.06, p=0.27, bandwidth vs latency: Pearson’s r=0.08, p=0.07, **Extended Figure 3B-C**).

To determine whether classifying the type of projection neuron improved functional categorization beyond what could be accomplished by reporting the laminar location of the cell body, we fitted a linear support vector machine (SVM) classifier model (Cortes & Vapnik 1995; Hastie et al. 2008) to both the CT (**Figure 4A**) and layer (**Figure 4B**) tuning parameters and tested, using cross-validation, how well either the correct cell-type or layer could be predicted. Classification was better than chance (50%) in either case, although classification errors were significantly lower based on projection cell type than for cortical layer (32% vs 23%, Bootstrapped statistical test, p<0.05, **Figure 4C**). This indicates that variance in sensory response properties can be better predicted by classification of neuron type, rather than just cortical layer.

**Figure 4.**
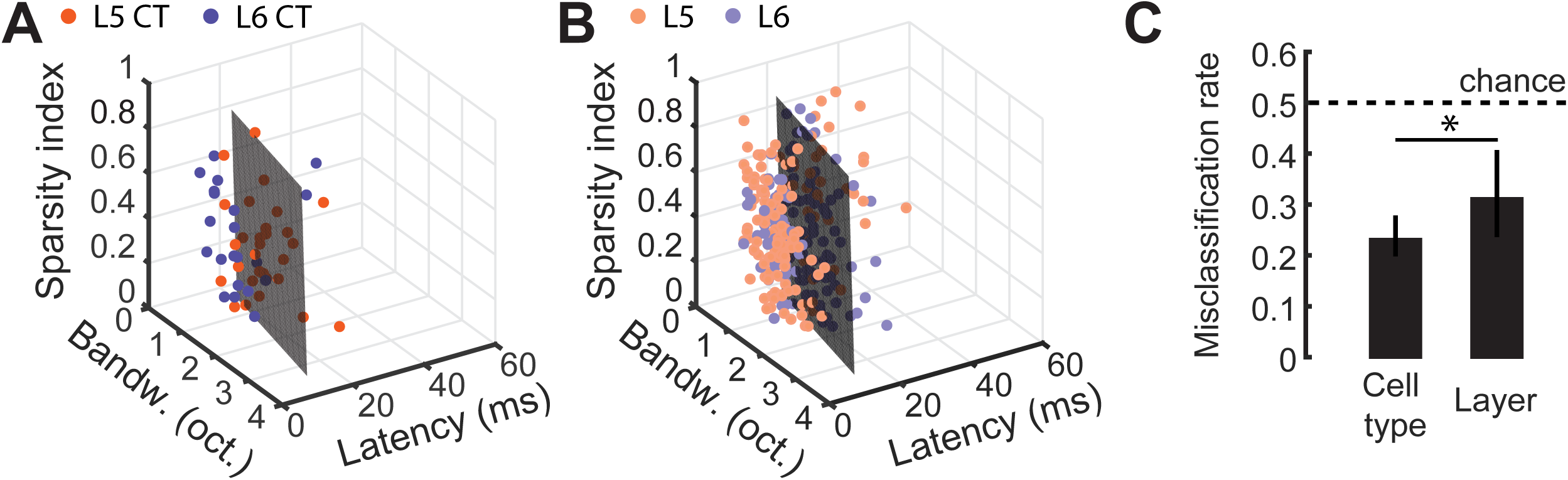
Predicting cell-type and layer from neural tuning parameters. **A:** Scatter plot of tuning parameters for both CT unit types, with the optimal SVM hyperplane. **B**: Scatter plot of tuning parameters for L5 and 6 unit types, with the optimal SVM hyperplane. **C**: Mean (± 95% bootstrapped CI) misclassification rates for cross-validated SVM fits to either cell-type or layer. Dotted line is chance, at 0.5. Asterisk denotes a statistically significant difference at p<0.05, using a Bootstrapped statistical test.

### Modeling the stimulus-response function of CT neurons

Fully characterizing the stimulus-response transformation of a neuron is an intractable problem as it would require knowing the neural responses to all possible stimuli. A common approach is to present a complex stimulus that spans a sizeable subset of the possible stimulus space, and to then relate the stimulus to the resultant neural response using mathematical models. We chose to do this by presenting 20 minutes of a dynamic random chord (DRC) stimulus (DeCharms et al. 1998; Linden 2003). Only neurons that were significantly driven by the DRC were considered for analysis (L5: n=372, L6: n=295, L5 CT: n=81, L6 CT: n=51). To mathematically describe the stimulus-response functions for both subtypes of CT neuron, we proceeded to fit a multilinear contextual gain field (CGF) model to the DRC responses (Ahrens et al. 2008; Williamson et al. 2016). Nonlinear contextual effects are prevalent within the auditory system, and this model is capable of capturing some of these known acoustic interactions within the stimulus that can lead to nonlinearities within the neural response (Brosch & Schreiner 1997; Wehr & Zador 2005; Kadia & Wang 2003; Sadagopan & Wang 2009; Phillips et al. 2017). We quantified predictive accuracy by computing the fraction of the stimulus-related variability could be captured by the model, and compared this to the accuracy achieved by a linear spectrotemporal receptive field (STRF) model (**Figure 5A**). The CGF model outperformed the linear STRF model on almost every cell (Wilcoxon Signed Rank test, L5: p<2×10^-52^, L6: p<5×10^-44^, L5 CT: p<2×10^-12^, L6CT: p<3×10^-7^, **Figure 5B**).

**Figure 5.**
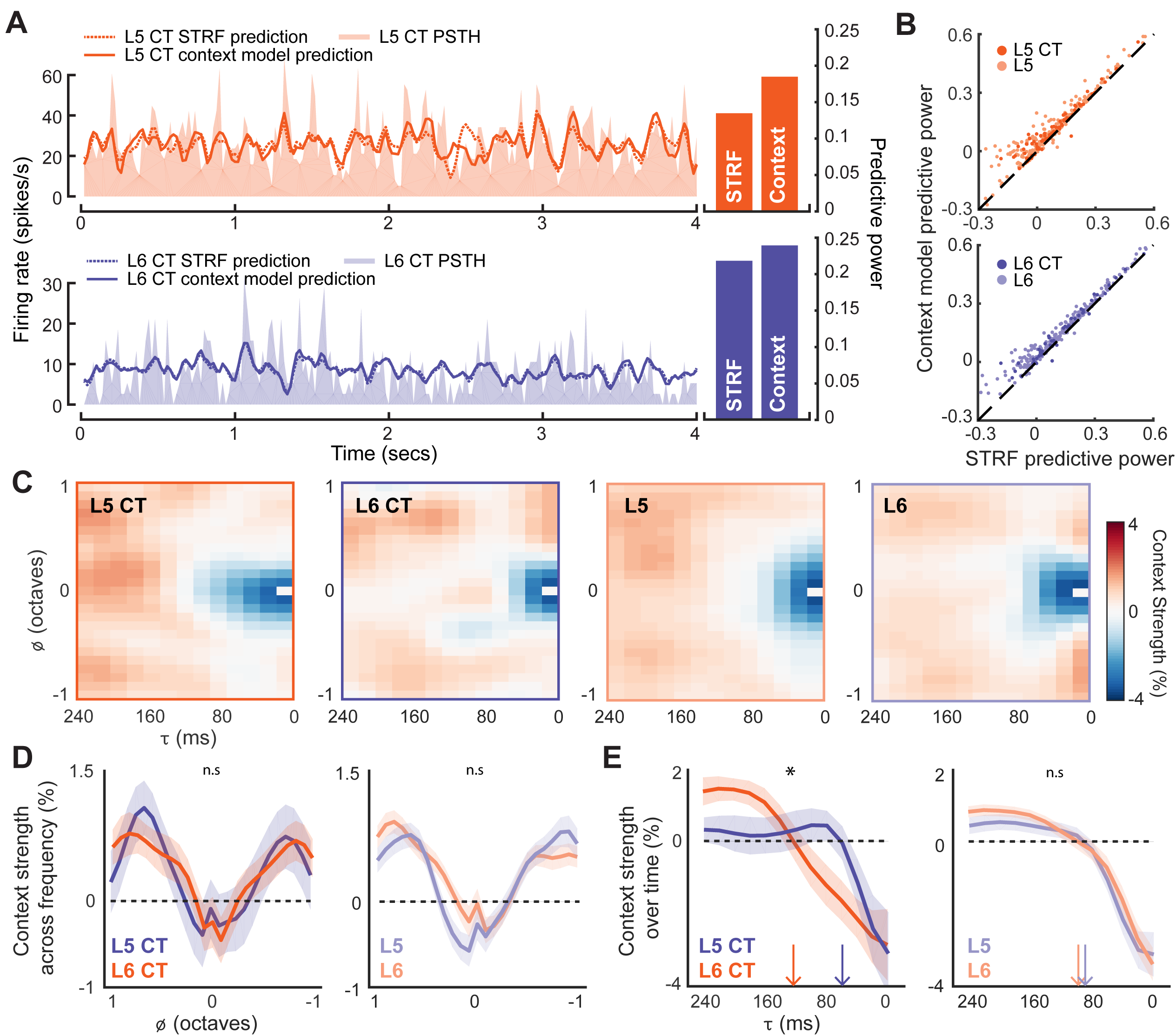
Modeling the stimulus-response _function ofL5 andL6 neurons. **A**: Snippets of DRC- evoked neural activity with corresponding model predictions for an example L5 CT (top) and L6 CT neuron (bottom). Predictive powers for the model fits are shown on the right. **B**: Scatter plots showing the relation between STRF and context model predictive powers for all recorded units. **C**: Mean contextual gain field (CGF) for each L5 and L6 unit subtype. **D**: Mean (±1 SEM) of an average across “for both CT subtypes (*left*) and layers (*right*). **E**: Mean (±1 SEM) of an average across a range of τ (between ±0.25 octaves) for both CT subtypes (*left*) and layers (*right*). Vertical arrows depict the extent of the suppressive timescale (the point at which the function crosses zero). Asterisk denotes a statistically significant difference at p<0.05, as determined by a mixed-effect ANOVA.

The CGF model combines two receptive field structures. The first is an STRF-like principal receptive field, while the second is the CGF. CGFs consistently featured a suppressive region centered at a zero frequency offset, indicating that preceding sound energy at a similar frequency tended to dampen the impact of a component sound. Regions of enhancement were also present at longer time delays near the preferred frequency, in addition to enhancement in spectral side lobes that overlapped in time with the inputs at the preferred frequency (**Figure 5C- D**) (Williamson et al. 2016). Nonlinear context effects did not substantially differ between L5 and L6 over frequency or time (mixed effect ANOVA, *τ* x cell-type interaction, F_(1,12)_ <0.5, p=0.95, Φ x cell-type interaction, F_(1,24)_<1.3, p=0.18, **Figure 5D-E**). Among the subset of CT units, however, we noted that the delayed suppressive region (τ) was significantly shorter in L6 CTs, suggesting that the nonlinear components of forward suppression operate on faster timescales in L6 CT neurons than L5 CT neurons (mixed effect ANOVA, *τ* x cell-type interaction, F_(1,12)_ >2.1, p=0.02, “x cell-type interaction, F_(1,24)_<0.3, p=0.99, Figure 5D-E).

### Local connectivity within deep-layer cortical networks

L6 CT neurons have previously been shown to exert a strong feedfoward influence onto the local cortical circuit through interactions with FS neurons (Bortone et al. 2014; Guo et al. 2017; Kim et al. 2014). However, the local influence of L5 CT neurons is unclear. To investigate feedforward excitation, we studied the effect of L5 CT or L6 CT activation on local populations of RS and FS neurons, while they were in a quiescent state lacking sensory stimulation. Optogenetic activation of L5 CT axons elicited weak, brief polysynaptic activation of neighboring RS and FS units in L5 (**Figure 6A**: *left*). By contrast, photoactivating the thalamic terminals of L6 CT units drove powerful, sustained and distributed activation of local FS and RS units (Figure 6A: *right*). We estimated the strength of local synaptic feedforward excitation from L5 CT and L6 CT neurons by quantifying RS and FS firing rates normalized to the peak of direct L5 CT or L6 CT activation. Whereas L5 CT evoked spiking rapidly extinguished in nearby RS and FS neurons, a single pulse of light to L6 CT axon terminals drove intense, prolonged polysynaptic activity in local RS and FS cells (Wilcoxon Rank Sum test; L5CT->FS (n=111) vs L6 CT->FS (n=73): p<4×10^-4^, L5CT->RS (n=652) vs L6 CT->RS (n=376): p<3×10^-25^, **Figure 6B-C**).

**Figure 6.**
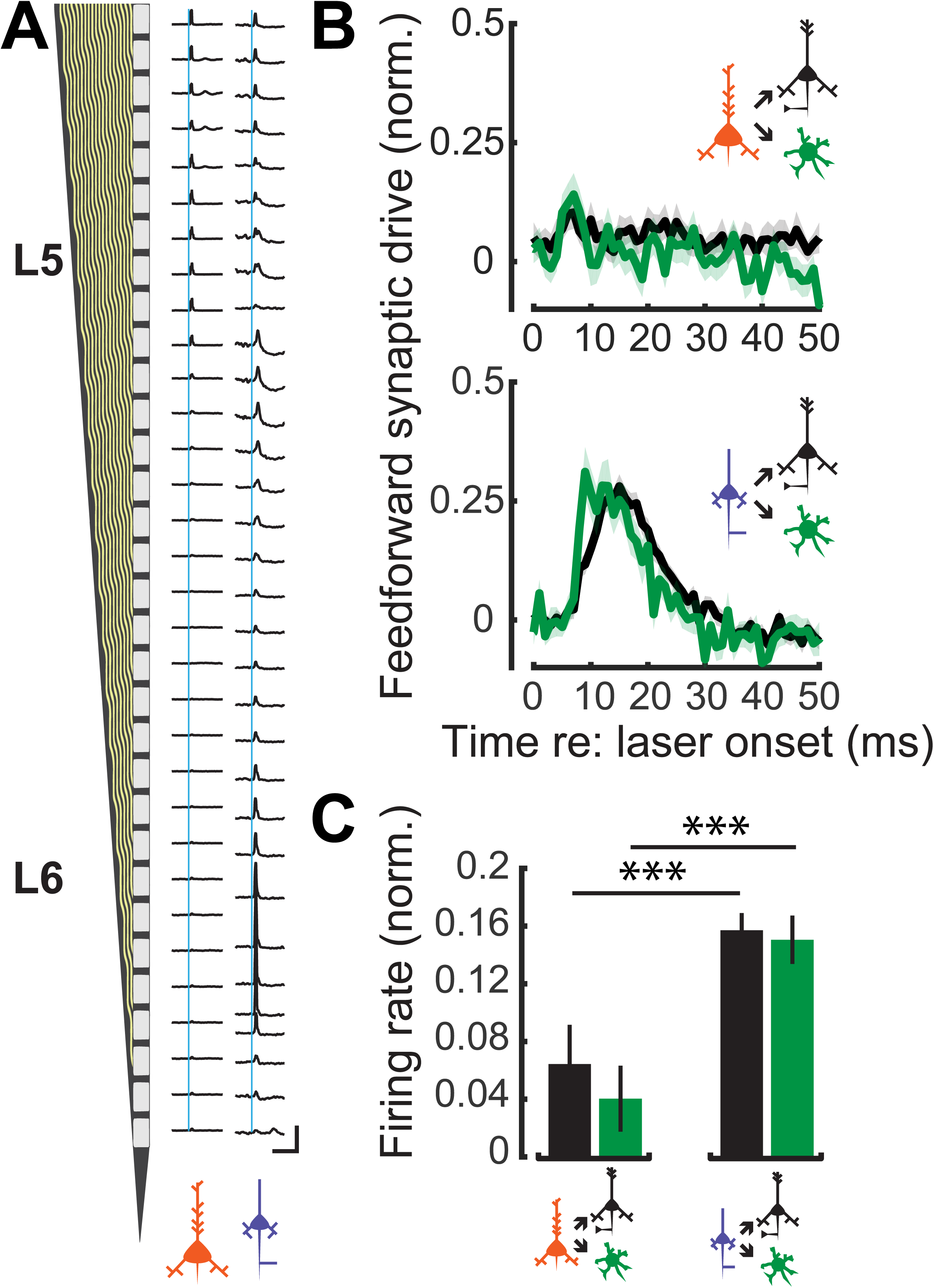
Feedforward connectivity within deep-layer cortical networks. **A**: Schematic of the 32- channel recording probe alongside PSTHs corresponding to each recording site. A 1 ms laser pulse (vertical blue lines) to the axon terminals for L5 CT (*left*) and L6 CT (*right*) recruits different patterns of spiking activity in representative deep layer recordings. Vertical scale bar = 200 sp/s; horizontal scale bar = 125 ms. **B**: Mean (±1 SEM) population normalized PSTHs for both RS and FS groups in response to either L5 CT activation (top) or L6 CT activation (bottom). **C**: Mean (±1 SEM) normalized firing rates in local RS and FS units, from the 5-25 ms epoch of the PSTHs shown in *B.* Asterisks denote statistically significant differences at p<0.005, as determined by the Wilcoxon Rank Sum test.

We then analyzed the temporal relationship in spiking patterns of both L5 CT and L6 CT neurons and the surrounding RS and FS population. We used a cross-covariance procedure to estimate the strength of temporal interactions during “steady-state” activation with the DRC stimulus (**Figure 7A-B**) (Atencio & Schreiner 2010; Rosenberg et al. 1989). The cross-covariance between RS neurons and both subtypes of CT neurons were relatively weak, although this interaction was stronger for L6 CT neurons (Wilcoxon Rank Sum test; L5CT<->RS (n=2553) vs L6CT<->RS (n=985): p<5×10^-8^, **Figure 7D-E**). Functional coupling with FS GABAergic interneurons, by contrast, was stronger overall and was particularly pronounced in L6 CT units (Wilcoxon Rank Sum test; L5CT<->FS (n=458) vs L6CT<->FS (n=209): p<5×10^-4^, Figure 7D-E).

**Figure 7:**
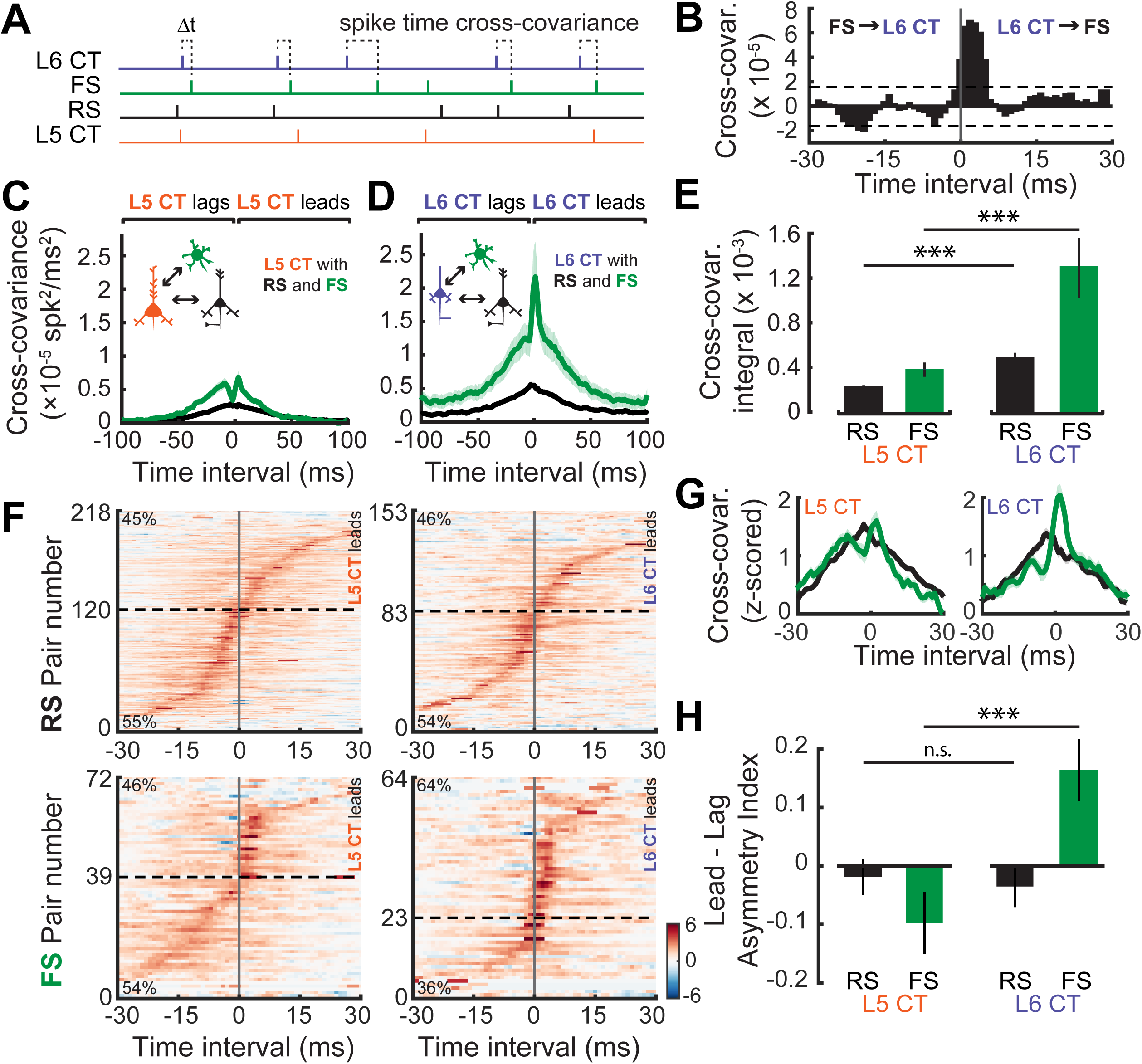
Temporal interactions within deep-layer cortical networks. **A**: Schematic illustrating the four recorded cell-types and the cross-covariance approach. **B**: An example cross-covariance function highlighting the temporal interaction between a L6 CT - FS pair. **C-D**: Mean (±1 SEM) cross-covariance functions of L5 CT (C) and L6 CT (D) neurons with RS and FS neuron types.**E**: Mean (±1 SEM) numerical integral of the cross-covariance functions in D-E. **F**: Significant cross-covariance functions were z-scored and sorted by peak timing. The proportion of pairs where the CT neuron either leads or lags are denoted in the top-left or bottom-left corners, respectively. Dashed horizontal line indicates the separation between leading and lagging pairs. Color scale bar denotes the z-scored cross-covariance. **G**: Marginal distributions of the z-scored cross-covariance functions shown in F. **H**: Mean (±1 SEM) asymmetry indexes between the positive and negative regions of the cross-covariance functions shown in F, where a value of 0 indicates equivalent regions of leading and lagging cross-covariance, negative values indicate a bias towards CT lagging spike times, and positive values indicate a bias towards CT leading spike times. Asterisks denote statistically significant differences at p<0.005, as determined by the Rank Sum test.

As a final step, we calculated the directionality of spike train covariance to test the idea that L5 CTs are integrators that pool inputs from local cells before broadcasting signals out of the cortex, whereas L6 CTs control columnar gain by driving local cell types. In this case, we hypothesized that L6 CT spikes would temporally lead spikes in local FS and RS cell types, whereas L5 CT spikes would lag behind local RS and FS spikes. To test this prediction, we first determined which spike train correlations were significant using a confidence bound, before z-scoring the cross-covariance functions and ordering them by the location of their peak (**Figure 7F**). The proportion of lead-preferring interactions was similar across L5 CT and RS/FS pairs, and L6 CT and RS pairs, at approximately 48%. By contrast, 65% of the L6 CT and FS pairs were lead-preferring indicating a preferential interaction between L6 CT neurons and neighboring FS GABA neurons. In addition to the temporal peak of the cross-covariance function, we also analyzed the amount of cross-covariance associated with each direction of interaction. We averaged both sides of the z-scored cross-covariance functions before computing a lead-lag asymmetry index by calculating (lead-lag)/(lead+lag) yielding a value whose distance from 0 indicates either lead-preferring (>0) or lag-preferring. We observed that L6 CT spikes tend to lead FS spikes, while the reverse is true for L5 CT spikes (Wilcoxon Rank Sum test; L5CT<->FS (n=56) vs L6CT<->FS (n=54), p<8×10^-4^, **Figure 7H**). These results echo previous reports that L6 CT neurons can dynamically regulate the gain of deep layer ACtx networks by driving networks of deep layer FS interneurons (Bortone et al. 2014; Guo et al. 2017).

## Discussion

Slice physiology studies have demonstrated that there is not a singular class of CT neuron. Rather, there are at least two parallel CT systems, one originating in L5, the other in L6, that differ systematically in their morphological, intrinsic, and synaptic properties (Diamond et al. 1969; Llano & Sherman 2009; Bajo et al. 1995; Rouiller & Welker 2000; Sherman & Guillery 2011; Ojima 1994; Bartlett et al. 2000; Andersen et al. 1980; Sherman 2016; Briggs et al. 2016). Technical limitations have made it challenging to extend these observations into the realm of sensory processing differences in intact, awake animals while still preserving single spike resolution at a cellular scale. Here, we overcame this technical limitation by leveraging recent advances in multi-channel electrophysiology and optogenetics to make targeted recordings from both L5 and L6 CT neurons in awake mice (Figure 2).

We showed that these pathways differ in both structure and function. In general, L5 CT neurons exhibited broader spectral tuning and lower lifetime sparseness, indicative of reduced stimulus selectivity and a broader response distribution (Figure 3). L5 CT neurons also exhibited a longer timecourse for suppressive stimulus interactions, suggesting an increased sensitivity to local acoustic context (Figure 5). Mechanistically, these differences in auditory tuning properties and nonlinear gain components likely reflect differences in thalamocortical input and local intracortical activity (Sun et al. 2013; Zhou et al. 2010). Analyses of functional connectivity showed that L6 CT neurons exerted strong feedforward synaptic drive onto local columnar neurons, while L5 CT drive was much weaker (Figure 6). L6 CT spiking activity also tended to lead the spiking of FS interneurons, suggesting an interaction with local inhibitory subnetworks (Figure 7). Taken together, these findings serve to highlight a functional dichotomy of sensory information processing between two cortical cell-types that both route information through the thalamus.

L5 CT neurons fall into the class of L5 pyramidal neurons referred to as intrinsic bursting (IB), based on their ability to fire bursts of action potentials in response to depolarizing current injection (Connors et al. 1982; McCormick et al. 1985). Previous work has investigated tuning bandwidths of such IB neurons and found them to be more broadly tuned than other neurons within L5 (Sun et al. 2013), a trend that is consistent with our observations. Moreover, L5 neurons also receive broad excitatory and narrow inhibitory input directly from the thalamus (Sun et al. 2013; Constantinople & Bruno 2013; Hooks et al. 2013), contributing to their broad tuning. Such input may lead to these neurons being more sensitive to nonlinear effects of acoustic context, consistent with our observations that the nonlinear components of forward suppression operate on longer timescales in L5 CT neurons. Recent work in the visual system has also described L5 CT neurons as forming a functional subnetwork within L5, that differs in connectivity and function from other L5 pyramidal neurons (Lur et al. 2016; Kim et al. 2015). The sparse tuning properties that we observed in L6 CT neurons echoes the sparse orientation tuning that has been observed in the visual system (Vélez-Fort et al. 2014).

Given that feedback connections grossly outnumber feedforward connections in the thalamus, these differences in response properties could influence the receptive fields of downstream thalamic neurons. Indeed, manipulating CT feedback can lead to modulation of receptive field properties in multiple sensory systems (Suga 2008; Alitto & Usrey 2003; He 1997; Wang et al. 2018; Temereanca & Simons 2004). The downstream targets of each CT cell- type also share some of the same differences in feature selectivity that we have reported. Previous work has shown that spectral tuning in MGBd is broader than in MGBv (Anderson & Linden 2011) reflecting the differences in tuning bandwidth we observed between L5 CT (primarily MGBd innervating) and L6 CT (primarily MGBv innervating). Broad spectral tuning is also shared with the external cortices of the IC (Ramachandran et al. 1999), where the L5 CT neurons also project. First-spike latencies in the MGBd are also almost double those found in MGBv, consistent with the differences we report between CT subtypes (Anderson & Linden 2011). This like-to-like functional organization is also reflected in higher-order feedback pathways in the visual system (Huh et al. 2018).

Ultimately, these findings shed light on how information contained in two primary output layers of the ACtx can be routed through the thalamus, to the rest of the brain. The L6 CT neurons propagate information that is sparse and more selective, but their anatomy places them in a position to modulate information both en route to the thalamus (through the TRN), but also locally through dense communication with the FS population. Such a circuit allows for rapid modulation of response gain and has led to speculation that L6 CT circuits play a crucial role in dynamically regulating stimulus salience according to internal state variables such as anticipation and attention (Guo et al. 2017; Homma et al. 2017).

Thanks in part to their elaborate dendritic structure, L5 CT neurons are able to integrate input from multiple ACtx layers and broadcast a dense, non-linear signal to multiple sub-cortical auditory stations. The large terminals deposited by L5 CT neurons in the MGBd will likely lead to increased spike reliability which will, in turn, ensure that a reliable signal is propagated from higher-order thalamus to higher-order areas of ACtx (Sherman 2012). The impact that these signals have on all target areas also need not be the same, as different downstream synaptic properties could lead to a postsynaptic variation in response.

Ascending sensory pathways are classically characterized as two streams: a lemniscal system for higher fidelity propagation of detailed stimulus information and a non-lemniscal system that captures contextual influences and internal state variables. Here we show that a division of labor between two parallel, complementary streams is maintained in the first-order component of the corticofugal system as well, underscoring a unifying theme in the organization of central sensory pathways.

## Materials and Methods

### Mice

All procedures were approved by the Massachusetts Eye and Ear Infirmary Animal Care and Use Committee and followed the guidelines established by the National Institute of Health for the care and use of laboratory animals. This study is based on data from 18 mice (aged 6-8 weeks, both male and female). All mice were maintained under a 12h/12h periodic light cycle with ad libitum access to food and water. Six C57Bl/6 mice were used for intersectional anatomy experiments (Figure 1). Two Ntsr1-Cre mice and 2 C57Bl/6 mice were used for anaesthetized pharmacology experiments (Figure 2). Six Ntsr1-Cre mice (B6. FVB(Cg)-Tg(Ntsr1-Cre) GN220Gsat/Mmcd) and 6 C57Bl/6 mice were used for awake electrophysiology experiments (Figures 3-8).

### Surgical Procedures

#### Virus-mediated gene delivery

Mice were anesthetized using isoflurane (4%). A surgical plane of anesthesia was maintained throughout the procedure using continuous infusion of isoflurane (1-2% in oxygen). A homoeothermic blanket system (Fine Science Tools) was used to maintain core body temperature at approximately 36.5°C. The surgical area was first shaved and cleaned with iodine and ethanol before being numbed with a subcutaneous injection of lidocaine (5 mg/mL). For viral delivery to the ACtx, an incision was made to the right side of the scalp to expose the skull around the caudal end of the temporal ridge. The temporalis muscle was then retracted and two burr holes (approximately 0.3 mm each) were made along the temporal ridge, spanning a region of 1.5 – 2.5 mm rostral to the lambdoid suture. A motorized stereotaxic injection system (Stoelting Co.) was used to inject 0.5 μl of either a non-specific ChR2 viral vector (AAV5-CamKIIa-hChR2[E123T/T159C]-eYFP, UNC Vector Core) or a Cre-dependent ChR2 viral vector (AAV5-EF1a-DIO-hChR2[E123T/T159C]-mCherry, UNC Vector Core) into each burr hole approximately 0.5 mm below the pial surface with an injection rate of 0.05 - 0.1 μl/min. Following the injection, the surgical area was sutured shut, antibiotic ointment was applied to the wound margin, and an analgesic was administered (Buprenex, 0.05 mg/kg). For intersectional anatomy experiments described in Figure 1, CAV2-Cre was injected into the IC via a small craniotomy atop the inferior colliculus (0.5 mm × 0.5 mm, medial-lateral x rostral-caudal, 0.25 mm caudal to the lambdoid suture, 1 mm lateral to midline). Following injections, the craniotomy was filled with antibiotic ointment (Bacitracin) and sealed with UV-cured cement. Neurophysiology experiments began 3-4 weeks following virus injection.

#### Implantation of optic fibers

Mice were brought to a surgical plane of anesthesia, using the same protocol for anesthesia and body temperature control described above. The dorsal surface of the skull was exposed, and the periosteum was thoroughly removed. If the animal was to undergo awake recordings, the skull was then prepared with 70% ethanol and etchant (C&B Metabond) before attaching a custom titanium head plate (eMachineShop) to the skull overlying bregma with dental cement (C&B Metabond). For L5 CT phototagging, a small craniotomy (0.5 mm × 0.5 mm, medial-lateral x rostral-caudal) was made (0.25 mm caudal to the lambdoid suture, 1 mm lateral to midline), to expose the IC. A ferrule and optic fiber assembly was positioned to rest atop the IC and then cemented to the surrounding skull (C&B Metabond). For mice undergoing L6 CT phototagging, a small craniotomy (0.5 × 0.5 mm, medial-lateral x rostral-caudal) was made under stereotaxic guidance, centered 2.75 mm lateral to midline and 2.75 mm caudal to bregma. A ferrule with a longer optic fiber (~3 mm) was implanted 2.7 mm below the brain surface to illuminate axons innervating the MGB. Once dry, the cement surrounding the fiber implant was painted black with nail polish to prevent light from escaping. Mice recovered in a warmed chamber and were housed individually.

### Acoustic and optogenetic stimulation

#### Acoustic stimulation

Stimuli were generated with a 24-bit digital-to-analog converter (National Instruments model PXI-4461) using custom scripts programmed in MATLAB (MathWorks) and LabVIEW (National Instruments). Stimuli were presented via a free-field electrostatic speaker (Tucker-Davis Technologies) facing the left (contralateral) ear. Free-field stimuli were calibrated before recording using a wide-band ultrasonic acoustic sensor (Knowles Acoustics, model SPM0204UD5).

#### Light delivery

Collimated blue light (488 nm) was generated by a diode laser (Omicron, LuxX) and delivered to the brain via an implanted multimode optic fiber coupled to the optical patch cable by a ceramic mating sleeve. Laser power through the optic fiber assembly was calibrated prior to implantation with a photodetector (Thorlabs).

### Neurophysiology

#### Awake, head-fixed preparation

Before the first awake recording session, mice were briefly anesthetized with isoflurane (4% induction, 1% maintenance) before using a scalpel to make a small craniotomy (0.5 mm x 0.5 mm, medial-lateral x rostral-caudal) atop the ACtx, at the caudal end of the right squamosal suture centered 1.5 mm rostral to the lambdoid suture. A small chamber was built around the craniotomy with UV-cured cement and a thin layer of silicon oil was applied to the surface of the brain. The mouse was then brought into the recording chamber and the head was immobilized by attaching the headplate to a rigid clamp (Altechna). We waited at least 30 minutes before starting neurophysiology recordings. All recordings were performed in a single-walled sound-attenuating booth lined with anechoic foam (Acoustic Systems).

At the end of each recording session, the cement chamber was flushed with saline, filled with antibiotic ointment (Bacitracin) and sealed with a cap of UV-cured cement. The chamber was removed and rebuilt under isoflurane anesthesia before each subsequent recording session. Typically, 3-5 recording sessions were performed on each animal over the course of 1 week.

#### Data acquisition and spike sorting

At the beginning of each session, a 32-channel, single-shank, silicon probe with 20 μm between contacts (NeuroNexus A32-Edge-5mm-20-177-Z32) was inserted into the ACtx perpendicular to the brain surface using a micromanipulator (Narishige) and a hydraulic microdrive (FHC). Once inserted, the brain was allowed to settle for 10-20 minutes before recording began. We ensured that recordings were made from the core fields of ACtx (either A1 or AAF) based on the tonotopic gradient, response latencies, and frequency response area shape (Guo et al. 2012).

Raw neural signals were digitized at 32-bit, 24.4kHz (RZ5 BioAmp Processor; TuckerDavis Technologies) and stored in binary format for offline analysis. The signal was bandpass filtered at 300-3000 Hz with a second-order Butterworth filter and movement artifacts were minimized through common-mode rejection. To extract local field potentials, the raw signals were notch filtered at 60 Hz and downsampled to 1000 Hz. Signals were then spatially smoothed across channels using a 5-point Hanning window. We computed the second spatial derivative of the local field potential to define the laminar current source density (CSD), which was used to assign each recorded unit to layer 5 or 6 (Guo et al., (2017), Extended Figure 2).

Spikes were sorted into single-unit clusters using Kilosort (Pachitariu et al. 2016). All data files from a given penetration were concatenated and sorted together so that the same unit could be tracked over the course of the experiment (~90 minutes), and to ensure that a unit drifting across contacts could be accurately assigned to the same cluster. Single-unit isolation was based on the presence of both a refractory period within the inter-spike-interval histogram, and an isolation distance (>10) indicating that single-unit clusters were well separated from the surrounding noise (Schmitzer-Torbert et al. 2005; Harris et al. 2016).

Once isolated, single units were classified as RS, FS, L5 CT or L6 CT. For the FS and RS classification, the trough-to-peak interval of the mean spike waveform formed a bimodal distribution, which we used to subdivide neurons with intervals exceeding 0.6 ms as RS, while neurons with intervals shorter than 0.5 ms were FS (Extended Figure 2C). We used an optogenetic approach to classify a subset of recorded units as either L5 CT or L6 CT. A 1 ms laser pulse (with power typically ranging from 10-50 mW) was presented between 250 and 1000 times at a rate of 4 Hz, to antidromically activate ACtx neurons. We then fit an analysis window around the laser-evoked spikes responsive neurons (operationally defined as recordings where the firing rate increased by at least 5 SD from the pre-laser baseline firing rate). We then computed the temporal jitter as the standard deviation of the first spike that occurred within the responsive window (Figure 2).

#### Anesthetized neuropharmacology experiments

Pharmacology control experiments were performed in an anesthetized preparation to ensure that the drug could be safely and accurately delivered to the middle layers of ACtx. Mice were anesthetized with ketamine (100 mg/kg) and xylazine (10 mg/kg). A homoeothermic blanket system (Fine Science Tools) was used to maintain core body temperature at approximately 36.5°C. A surgical plane of anesthesia was maintained throughout the procedure with supplements of ketamine (50 mg/kg) as needed. An ACtx craniotomy was made as described earlier. In order to block synaptic transmission, we prepared a 1.0 mM NBQX solution by dissolving NBQX (Sigma) into artificial cerebrospinal fluid (ACSF, Harvard Apparatus). After baseline extracellular recordings with a 16 channel silicon probe (NeuroNexus A1×16-100-177-3mm), the NBQX solution was injected approximately 500 μm below the pial surface with an injection rate of 0.05 – 0.1 μl/min. We then proceeded with optogenetic stimulation experiments after first confirming that sound-evoked local field potential (LFP) responses were qualitatively eliminated.

### Electrophysiological data analyses

#### Frequency response areas

FRAs were delineated using pseudorandomly presented pure tones (50 ms duration, 4 ms raised cosine onset/offset ramps) of variable frequency (4-64 kHz in 0.1 octave increments) and level (0-60 dB in 5 dB increments). Each pure tone was repeated 2 times and responses to each iteration were averaged. Spikes were collected from the tone-driven portion of the PSTH. First-spike latencies were defined as the time at which the firing rate began to exceed the spontaneous rate by 3 SD. Bandwidths were defined as the FRA width, 20 dB above threshold. An FRA was considered well-defined if it had a d’ value of greater than 3.5. Additional details on FRA boundary determination and d’ computation have been described previously (Guo et al. 2012).

#### Sparseness

To probe the sparsity of our neuronal population, we generated a bank of random stimuli (Figure 3E), in which each stimulus consisted of a noise token that varied across four acoustic dimensions (center frequency (4-64 kHz, 0.1 octave steps), spectral bandwidth (0-1.5 octaves, 0.1 octave steps), level (0-60 dB SPL, 10 dB SPL steps), and sinusoidal amplitude modulation rate (0-70 Hz, 10 Hz steps)). This bank of stimuli was repeated 3-10 times, and the responses averaged.

We used a common measure of sparseness (Chambers et al. 2014; Vinje & Gallant 2000; Rolls & Tovee 1995), defined as

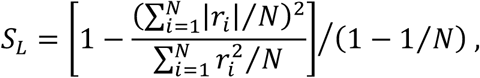

where *r_i_* represents firing rates, and i indexes time. This index is defined between zero (less sparse) and one (more sparse), and depends on the shape of the response distribution *p*(*r*).

#### Support vector machine classification

To determine how well tuning parameters could be used to correctly decode either cell-type or layer, we used a binary support vector machine classifier with a linear kernel. We fitted the classifier model to a data matrix consisting of N observations of 3 tuning parameters (bandwidth, latency, and sparsity). Each observation was associated with either a positive or negative class (which was used to indicate either cell-type or layer). Ten-fold cross-validation was then used to compute a misclassification rate. The SVM training and cross-validation procedure was carried out in MATLAB using the ‘fitcsvm’, ‘crossval’, and ‘kfoldLoss’ functions. Uncertainty in the misclassification rates was then quantified using a bootstrapping procedure to compute the 95% confidence intervals for both cell-type and layer. A statistically significant difference between groups was established if the classification accuracy of one group fell outside the bootstrapped 95% confidence interval of the other.

#### Statistical models of complex sound responses

We presented a 4-64 kHz dynamic random chord (DRC) stimulus (Linden, (2003)). This spectrotemporally rich stimulus is clocked such that every 20 ms a combination of 20 ms cosine-gated tone pulses with randomly chosen frequencies and intensities is generated. Frequencies were discretized into 1/12 octave bins spanning 4-64 kHz, and levels were discretized into 5 dB SPL bins spanning 25-70 dB SPL. The number of tones in each 20 ms chord was random, with an average density of 2 tone pulse per octave. A 1 min DRC was generated, and this was repeated for 20 trials, allowing for 20 min of continuous sound presentation.

We binned our PSTHs into 20 ms bins (to match the temporal resolution of the DRC) to yield a set of *N* response vectors 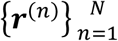, where *N* is the number of DRC repeats. We fitted both linear STRF and multilinear context models to the data. For a stimulus *s*, the linear STRF model is defined as

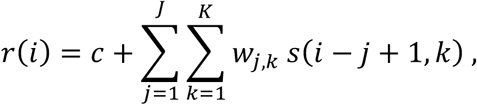

yielding a prediction of a neural firing rate *ř* at time 9, with *j* indexing time and *k* indexing frequency. This model can be extended to include the effects of acoustic context as

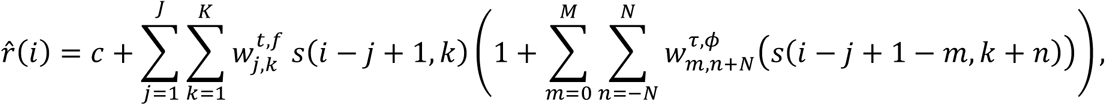

with m indexing *relative* time and n indexing *relative* frequency. The model consists of a linear component, with a principal receptive field (PRF; analogous to an STRF) denoted by w^*tf*^, and a contextual gain field (CGF) denoted by w^*τφ*^, which acts to multiplicatively modulate the stimulus spectrogram prior to spectrotemporal summation by the PRF.

Estimation of the STRF was carried out using the automatic smoothness determination (ASD) algorithm (Sahani & Linden 2003a; Meyer et al. 2017). This approach uses regularized linear regression with a spectrotemporal smoothness constraint which is optimized separately for each recording. Estimation of the PRF and CGF in the multilinear model was carried out using

ASD in an iterative alternating least squares (ALS) algorithm (Williamson et al. 2016; Ahrens et al. 2008).

Following Sahani & Linden, (2003b), we then define an estimator for the stimulus-dependent variability within the response, the signal power 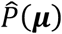

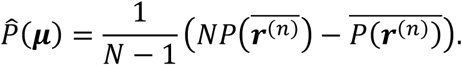

An estimate of the stimulus-independent trial-to-trial variability, the noise power, was obtained by subtracting this expression from 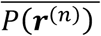. The DRC stimulus drove comparable spiking variability in both CT cell types (two-sample t-test, p=0.31,Extended Figure 5A-B). Noise power was also not different, (two-sample t-test, p=0.73, Extended Figure 5C) indicating that trial-to-trial variability was similar between groups.

The predictive performance of both models was evaluated using “predictive power”, an explainable variance metric that is normalized by the signal power (Sahani & Linden 2003b). It is therefore a measure of performance that explicitly takes trial-to-trial variability into account, and provides a means to quantify how much stimulus-related variability can be explained by a model. Its expected value for a perfect model is 1, and for a model predicting only the mean firing rate it is 0. Generalization performance of the models was assessed using 10-fold cross validation.

As described previously, predictive performance on both training and test data depended systematically on the amount of trial-to-trial variability in the recording (Extended Figure 5D-E) (Sahani & Linden 2003b; Williamson et al. 2016; Meyer et al. 2017; Ahrens et al. 2008; Englitz et al. 2010). Following these previous studies, we obtained population-level predictive performance estimates by extrapolating to the “zero-noise” limit, effectively averaging across the population while discounting the variable impact of noise on each neuron. These extrapolated limits bracket the true average predictive power of the model.

To further investigate the relevance of estimated CGF structure, we quantified the primary modes of variability around the mean using principal components analysis (PCA). In all groups, the scatter around the mean was concentrated in the first two or three principal components (PCs). The first three PCs were able to account for almost 60% of the total variability (Extended Figure 5F). The structure present within these PCs reflected the consistent features observed in the population CGFs. Namely, the dominant effect in the first PC was to modulate the overall depth of simultaneous/near-simultaneous enhancement, and in the second PC was to modulate the overall depth of delayed suppression (Extended Figure 5G).

#### Functional connectivity

Functional connectivity was assessed using cross-covariance methods (Rosenberg et al. 1989). For all pairs of spike trains, we followed Atencio & Schreiner, (2010) and first computed a cross-correlation function

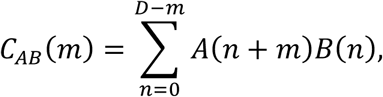

where b and B are 1 ms binned spike trains and D is the total duration. This cross-correlation function then leads to an unbiased estimate of the second-order cross-product density, *P_AB(m)_* as

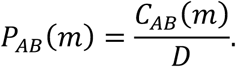

Defining the mean intensities of each spike train as 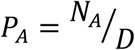 and 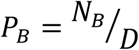, where *N_A_* and N_B_ are the total number of spikes in trains b and B, respectively, then leads to the crosscovariance function

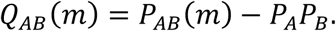

### Anatomy

Mice were deeply anesthetized with ketamine and transcardially perfused with 4% paraformaldehyde in 0.01M phosphate buffered saline. The brains were extracted and stored in 4% paraformaldehyde for 12 hours before transferring to cryoprotectant (30% sucrose) for 48 hours. Sections (40 μm) were cut using a cryostat (Leica CM3050S), mounted on glass slides and coverslipped (Vectashield). Fluorescence images were obtained with a confocal microscope (Leica).

### Statistical analyses

All statistical analysis was performed with MATLAB (Mathworks). Data are reported as mean ± SEM unless otherwise stated. Non-parametric statistical tests were used where data samples did not meet the assumptions of parametric statistical tests. In cases where the same data sample was used for multiple comparisons, all p-values remained significant after correction using the Benjamini-Hochberg procedure.

---

**Extended Figure 2.**
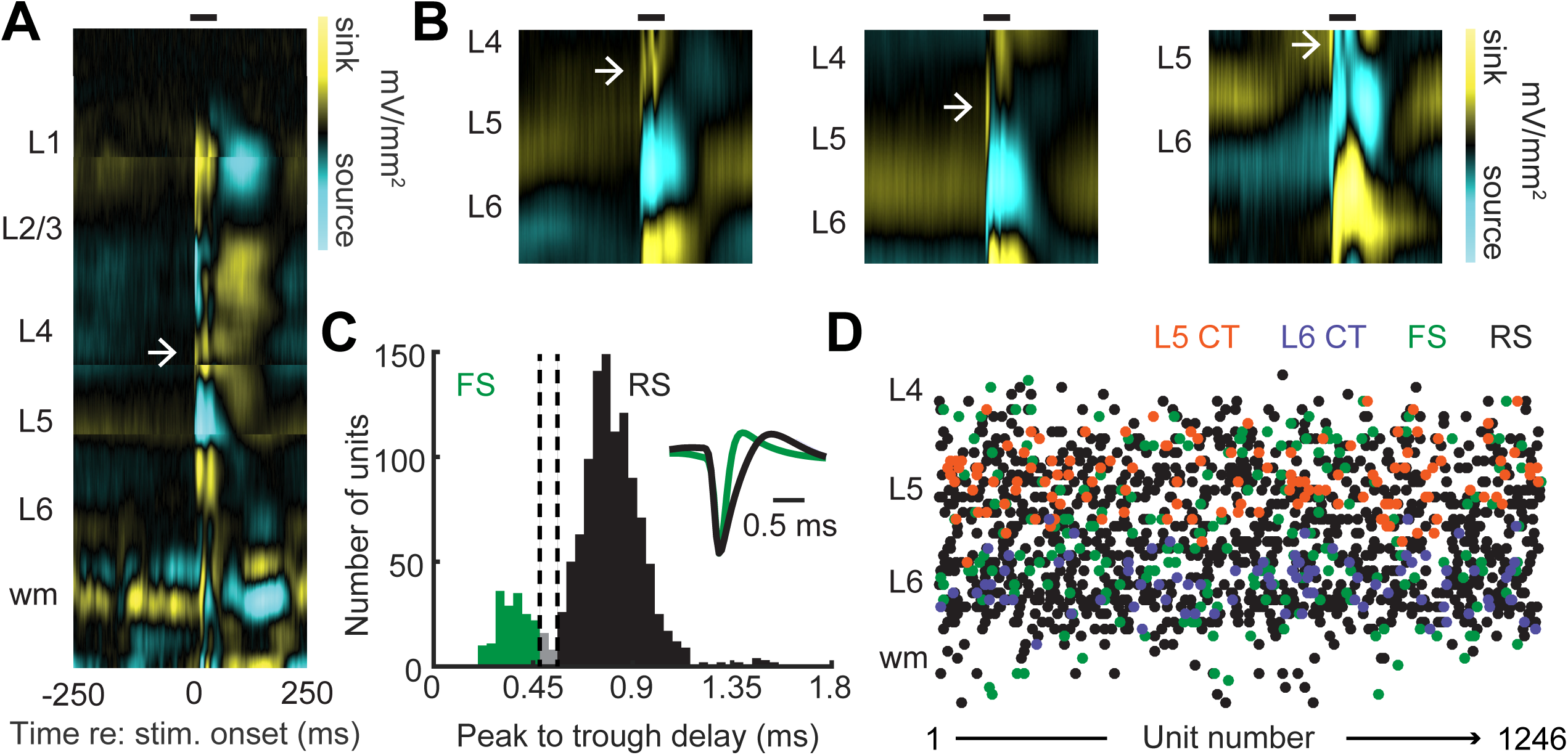
**A**: Four noise-evoked current source density plots (CSD) from recordings at different cortical depths. The CSD’s have been positioned to resemble the global CSD patterning of sinks and sources over the entire cortical column. Black horizontal line depicts timing of a 50 ms noise burst. White arrow depicts the location of the earliest sink, used to approximate the transition between ACtx L4 and 5. **B**: Three exemplar CSDs recorded from different mice. The 32-channel probe spans 610 μm and primarily covers ACtx L5 and 6. **C**: Histogram of spike waveform peak to trough delay values. Dashed vertical lines depict cutoffs used to classify units as RS (peak-to-trough delay < 0.5 ms) or FS (peak-to-trough delay > 0.6 ms). **D**: Each recorded neuron was assigned an approximate layer based on the pattern of sinks and sources in the CSD.

**Extended Figure 3.**
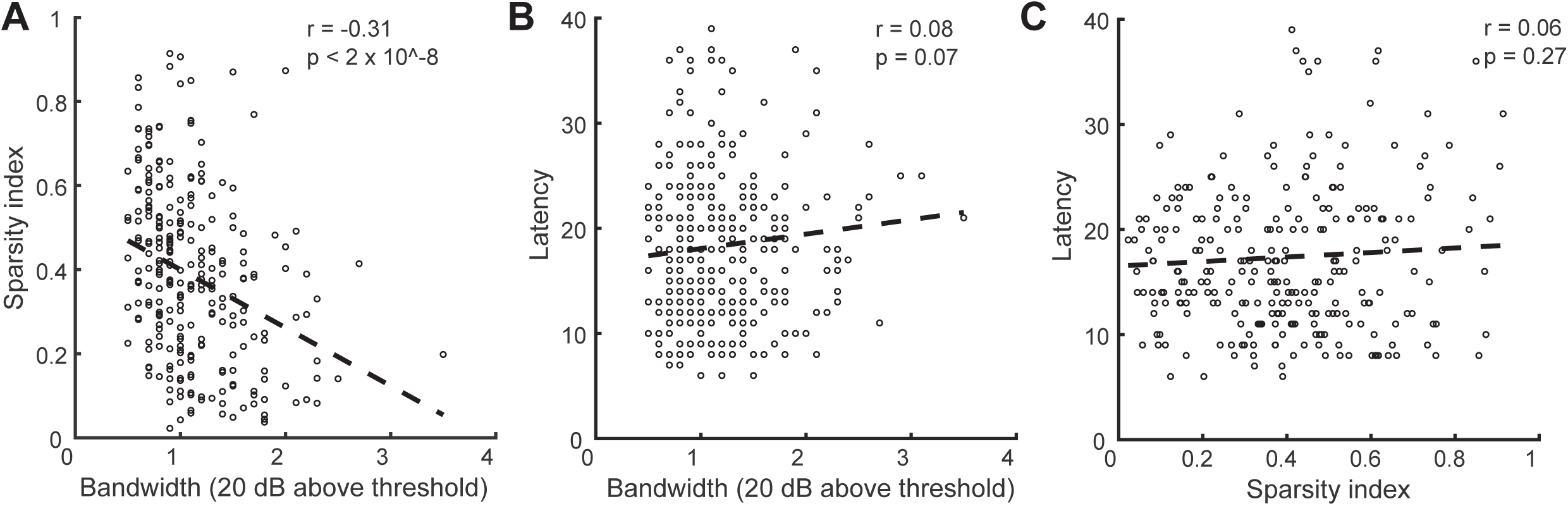
**A**: Significant correlation between sparsity and bandwidth parameters. Dotted line represents a linear fit to the data. **B**: No significant correlation between latency and bandwidth. **C**: No significant correlation between latency and sparsity.

**Extended Figure 5.**
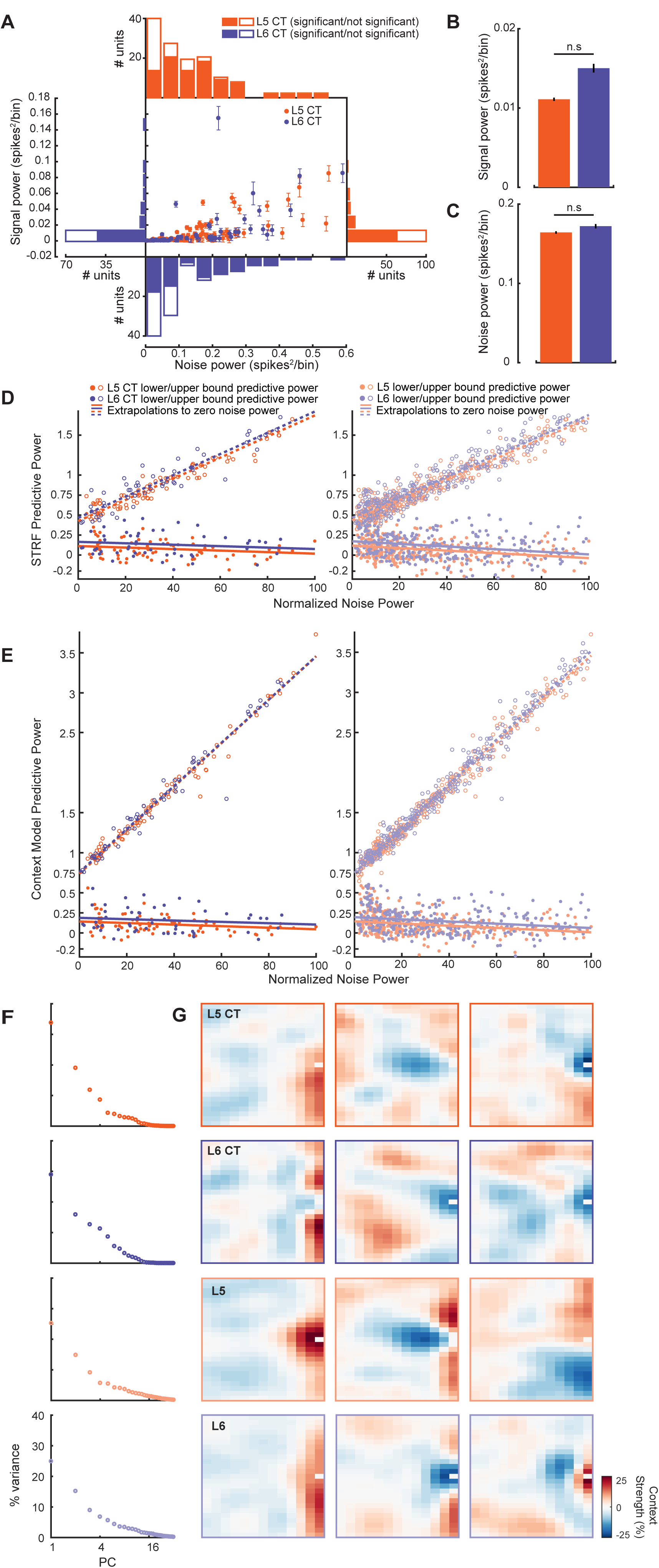
**A**: Scatter plot of signal and noise powers for both CT subtypes. Histograms show the marginal distributions along the corresponding axes. Filled/non filled bars represent neurons that had signal powers significantly greater than 0. **B**: Mean (±1 SEM) signal powers. **C**: Mean (±1 SEM) noise powers. **D**: Upper/lower bound STRF (*left*) and CGF (*right*) predictive powers plotted as a function of normalized noise power for both CT cell-types. Dotted/filled lines represent upper/lower bound extrapolations to the zero-noise condition. **E**: Upper/lower bound STRF (*left*) and CGF (*right*) predictive powers plotted as a function of normalized noise power for L5 and 6. Dotted/filled lines represent upper/lower bound extrapolations to the zero-noise condition. **F**: The absolute variance captured by each of the first 32 principal components, from a PCA analysis on the CGF populations. In each case, the first two or three principal components account for a large proportion of the variance. **G:** The 3 leading principal components (ordered left to right) for each CGF population. In all cases, the first two principal components indicate the primary sources of variability to be modulation of both delayed suppression, and facilitation of tones at short delays and nearby frequencies.

## Acknowledgements

We thank K. Hancock for support with data collection software, W. Guo for contributing to electrophysiological data analysis, and L. Sheets for guidance with pharmacology. We thank M. Pachitariu for both providing and assisting with the use of Kilosort. This work was supported by NIH grants R01 DC017078 (to DBP) and F32 DC01536 (to RSW).

**Author contributions:** RSW and DP conceptualized all experiments. RSW collected and analyzed all data. RSW and DP prepared figures and wrote the manuscript.

